# Vapor pressure deficit dominates the spatiotemporal variations in ecosystem photosynthetic quantum yield

**DOI:** 10.1101/2024.09.17.613385

**Authors:** Liyao Yu, Xiangzhong Luo, Ruiying Zhao, Tin W. Satriawan, Jiaqi Tian

## Abstract

- The quantum yield (*α*) of photosynthesis represents the maximum efficiency of light use as indicated by the initial slope of photosynthetic light response curves. Understanding *α* is crucial for accurate modeling of photosynthesis and terrestrial carbon cycle. Despite its importance, the spatial and temporal variations in *α* at large scales remain largely elusive.
- We leveraged long-term eddy-covariance observations from 90 sites globally and examined the spatiotemporal variations in *α* due to climatic drivers, using statistical and machine learning approaches.
- We found significant spatial variability in *α* across and within biomes, primarily driven by atmospheric vapor pressure deficit and soil moisture variations. Meanwhile, the temporal changes in *α* are mainly driven by the negative effect of vapor pressure deficit, which weakens the positive effects of elevated CO_2_ and leaf area index.
- Our results highlight the dominant role of vapor pressure deficit in controlling the spatiotemporal variations of *α* as well as the unneglectable impacts of soil water content, CO_2_, and leaf area on *α*. Those new results provide insights for improving the representation of *α* in ecosystem photosynthesis models.

## Introduction

Terrestrial photosynthesis is the largest component in global carbon cycle (Friedlingstein et al., 2023). Accurate estimation of its magnitude is crucial to simulate the dynamics of the global carbon cycle and project future climate change (Beer et al., 2010; Keenan & Williams, 2018). However, model estimations of terrestrial photosynthesis vary from 110 to 170 GtC year^−1^ (Anav et al., 2015; Beer et al., 2010; Piao et al., 2013; Ryu et al., 2019), indicating a large uncertainty that needs to be constrained. One commonly used model for ecosystem photosynthesis (gross primary product or GPP) simulation is the light use efficiency (LUE) model (Running et al., 2004; Xiao et al., 2004; Zhao et al., 2005), where GPP is the product of a maximum LUE (LUE_max_, or equivalently referred to as photosynthetic quantum yield *α*) with scalars of climate variables and the absorbed photosynthetically active radiation (APAR). Based on the structure of the LUE model and previous studies, the variation in GPP is highly dependent on the variation in *α* (Garbulsky et al., 2014).

At the leaf level, *α* has been well established and quantified as the initial slope from the non-linear light response curve of GPP (Johnson & Goody, 2011). In a theoretically non-stress condition, *α* should be 0.125 mol CO_2_ mol photons^−1^, as 1 mol CO_2_ assimilation requires at least 8 mol photons absorbed and passed through the electron chain (Walker, 1992). A few studies confirmed that the *α* values under non-stressed conditions (0.10–0.11 mol CO_2_ mol photons^−1^) in the labs are consistent with the value above and do not exhibit a variation among species or the environmental conditions at their habitats (Björkman & Demmig, 1987; Ehleringer & Björkman, 1977; Long et al., 1993). However, actual leaf- or individual-level *α* values are often lower than the above mentioned values due to abiotic stress conditions such as cold stress (Rogers et al., 2019; Xu et al., 2022) and drought (Arslan et al., 2023; Fang et al., 2023), light quality (Urban et al., 2007), or canopy density (Luo et al., 2000).

At the ecosystem level, some studies found that *α* exhibits large spatial variability (Rogers et al., 2017) and is sensitive to changes in water availability and temperature (Coops et al., 2010; Medlyn, 1998; Sandoval et al., 2023). Although models have adopted biome-specific *α* values (Arslan et al., 2023; Fang et al., 2023; Luo et al., 2000; Rogers et al., 2019; Urban et al., 2007; Xu et al., 2022) and considered the impacts of spatial variation in *α* to a certain degree, this empirical consideration neglects the impacts of environment gradient on *α* within a biome or the temporal changes in *α* due to climate change. In particular, the impact of elevated CO_2_ on ecosystem *α* has been heavily studied and is expected to cause an increase in ecosystem *α* due to the CO_2_ fertilization effect (De Kauwe et al., 2016; Keenan et al., 2023; Kolby Smith et al., 2016; Smith et al., 2020). Temperature changes are also likely to influence leaf photosynthetic capacity through acclimation and adaptation (Kumarathunge et al., 2019), which in turn induce changes in ecosystem *α.* In addition to the impact of long-term CO_2_ elevation (and its associated increase in ecosystem leaf area index, Zhu et al. (2016)) and warming on the trend of *α*, we also hypothesize that variations in water availability (i.e., vapor pressure deficit and soil moisture), light quality (i.e., diffuse light fraction), and leaf area index (reflecting vegetation density) might affect the variability in *α*, based on previously mentioned leaf-level knowledge.

To elucidate the environmental drivers on the spatial and temporal variations in ecosystem *α,* here we leveraged long-term (> 10 years) eddy covariance observations at 90 sites and established light response curves of net ecosystem exchange to extract *α* (see Methods). We aim to disentangle the dependence of *α* on a wide range of environmental variables (i.e., air temperature, CO_2_ concentration, soil moisture, vapor pressure deficit, diffuse light fraction, and leaf area index) and quantified their respective contributions to the spatial and temporal variations in *α* using statistical and machine learning approaches.

## Materials and Methods

### Eddy covariance and climate datasets

We used three standardised flux data products in our study: AmeriFlux Fluxnet (https://ameriflux.lbl.gov/data/flux-data-products/), ICOS Archive (https://www.icos-cp.eu/data-products), and FLUXNET2015 (https://fluxnet.org/), which were all processed by the FLUXNET2015 pipeline (Pastorello et al., 2020). We only considered sites with more than 10 years of eddy covariance and meteorological observations. Based on this criterion, 111 sites covering a wide range of 11 biomes were identified. We excluded US-Ne1 due to some questionable records of CO_2_ concentrations (> 700 μmol mol^−1^). We extracted the start and end of growing season at each site using R ‘*phenofit’* package (Kong et al., 2022) with a threshold of 20% of maximum daily GPP (Körner et al., 2023) and excluded several tropical and Arctic sites because lack of reasonable extracted growing seasons in these sites. We obtained 90 sites and summarised the site information in Table S1. Since the observations of photosynthetic photon flux density (PPFD) are unavailable at many sites, we used the shortwave solar radiation (SW) measured at each site as a proxy of PPFD, which is often about half of SW (Liang & Wang, 2020). We retrieved half-monthly LAI from the Global Inventory Modelling and Mapping Studies LAI product (GIMMS LAI4g v1.2, Cao et al. (2023)), root-zone soil water content (SWC) and potential evapotranspiration (PET) from the Global Land Evaporation Amsterdam Model (GLEAM) v3 dataset (Martens et al., 2017). We estimated the cloudiness index as 1 – SW_IN_ / SW_POT_ (Turner et al., 2003), in which SW_IN_ is the incident SW and SW_POT_ is the modelled potential SW at the land surface under a clear sky. Both SW_IN_ and SW_POT_ were provided in the flux datasets. We then used the cloudiness index as a proxy for the fraction of diffuse light (DF) at the site, with greater cloudiness indicating a greater DF. We calculated the aridity index (AI) as PET/P, where P is precipitation provided by the flux datasets. A higher AI value indicates more arid conditions.

### Derivation of *α* and LUE from eddy covariance observations

We fitted the response of non-gap-filled net ecosystem exchange (NEE) to SW with a rectangular hyperbola equation to derive *α* in a short duration (2–14 days) (Luo & Keenan, 2020):

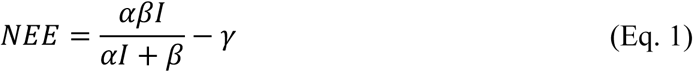

where *I* is the incident shortwave radiation (SW_IN_, μmol m^−2^ s^−1^), *β* is the maximum ecosystem photosynthetic rate (GPP_max_, μmol m^−2^ s^−1^), and *γ* is the ecosystem respiration rate (ER, μmol m^−2^ s^−1^). We derived GPP and ER using the daytime flux partitioning method (Lasslop et al., 2010). The fitting of the light response curve was embedded in the daytime flux partitioning method, and we used it to obtain daily *α* during the growing season, where we assumed *α* in each time window of 2–14 days remained the same. We provided an example of the curve fitting in Fig. 1. We then excluded physiologically unreasonable *α* (≥ 0.22 mol CO_2_ mol photons^−1^, twice of leaf-level maximum 0.11 mol CO_2_ mol photons^−1^, as SW_IN_ = 0.5 × PPFD) and estimated the growing-season average *α* based on the daily *α*.

**Fig. 1.**
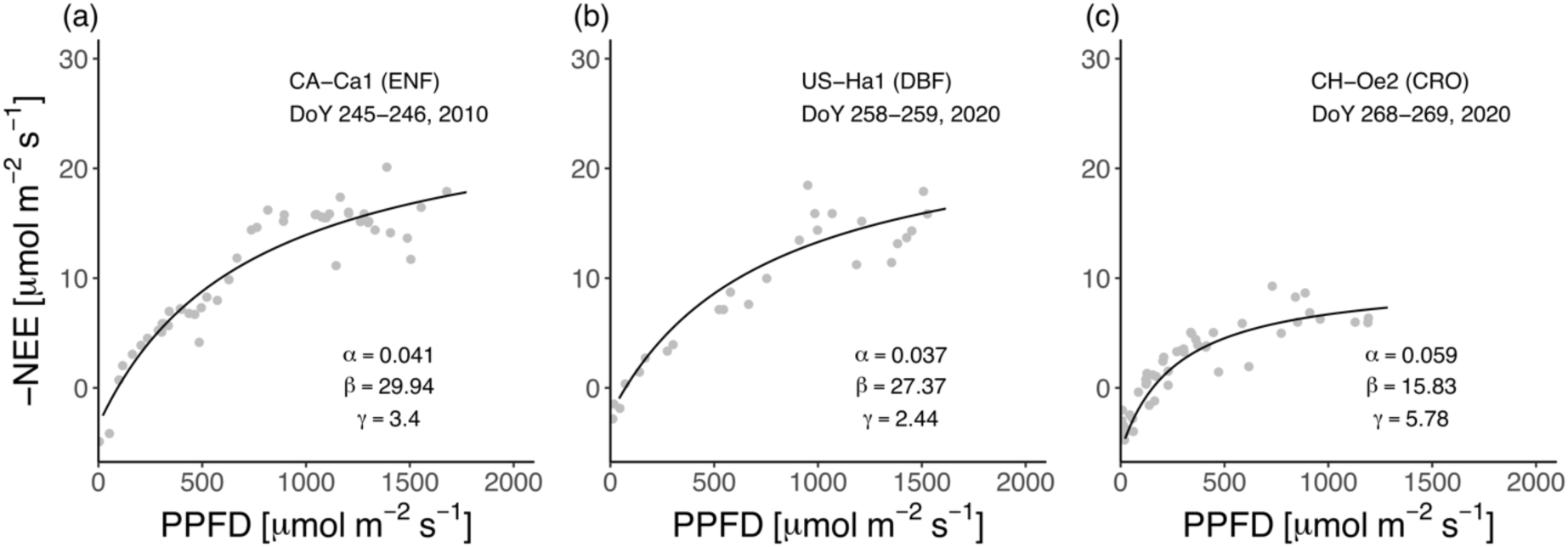
Examples of light response curves fitted to eddy covariance observations at three sites: (a) CA-Ca1, (b) US-Ha1, and (c) CH-Oe2. Grey dots indicate half-hourly or hourly observations of incident photosynthetic photon flux density (PPFD) and negative net ecosystem exchange (NEE) in a two-day window. The black lines indicate the fitted light response curve using the daytime partitioning approach (see Methods). We extracted the quantum yield (*α*), the maximum gross primary productivity (*β*), and the mean daytime ecosystem respiration rate (*γ*) from the fitted light response curves.

To facilitate the comparison between *α* and LUE, we also calculated half-monthly LUE as GPP/(fAPAR×*I*), where fAPAR is the fraction of absorbed photosynthetically active radiation inferred as 1−*e*^(−0.5×LAI)^. We then excluded physiologically unreasonable LUE values (≥ 0.22 mol CO_2_ mol photons^−1^, the same as above) and aggregated the LUE of each site to its annual growing season average.

### Explaining the spatial variation in *α*

To study the spatial variation in *α*, we investigated the variation in the site-average *α* across sites and used statistical methods to analyze the drivers of the cross-site variation. Prior to the analysis, we standardized all variables (i.e., *α*, climate, and LAI) as site means across all years. First, we built a linear model as:

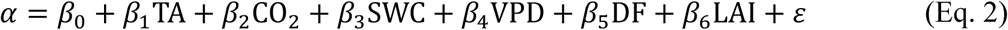

where *β*_0_ is an intercept, *β*_1_–*β*_6_ are the coefficient for the variables respectively, and *ε* is the unexplained error by the model. Since all the variables were standardized to a comparable scale, the coefficients (*β*_1_–*β*_6_) could reflect the relative strength to which the climate influences *α*. We then leveraged extended linear models, ridge regression and *lasso*, to analyse the drivers of spatial variations in *α*. These models are utilized to address the multicollinearity among climatic variables, such as the confounding effects between TA and VPD and between VPD and DF (Fig. S1). Ridge regression (Hoerl & Kennard, 2000) is a shrinkage method that applies a tuning parameter (λ) as L2 regularization to the original least-squares fitting and involves a penalty to the regression coefficients to reduce variance. This method has been widely applied to ecological studies to improve the reliability of the sensitivity analysis under multicollinearity (Peng et al., 2013; Zhong et al., 2023). We determined the best λ in the ridge regression model with 5-fold cross-validation using the R ‘*glmnet*’ package (Friedman et al., 2010), which minimizes the following objective:

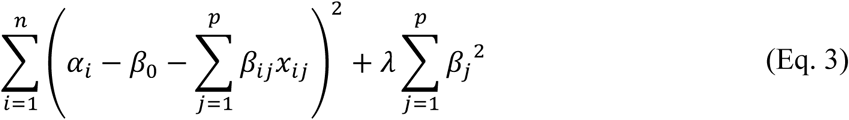

where *n* denotes sample size, *p* denotes and number of variables, and *x_j_* (1 ≤*j* ≤6) represents the variables. We also applied another widely used shrinkage method *lasso* (Tibshirani, 1996) (L1 regularization) with similar λ tuning approach and reported the averaged coefficients from the ridge regression and *lasso*.

### Explaining the temporal variations in *α*

For the temporal variation analysis, we first obtained the temporal anomalies of climate variables at each site by removing the site mean value from site-year value. This step is meant to eliminate the influence of spatial variation in climate. For instance, we calculated the anomaly of TA at a site in year *i* (TA_anom, *i*_) as TA*_i_*−TA_mean_, where TA*_i_* is the annual TA in year *i,* and TA_mean_ is the mean TA at that site across all years. We showed the correlation matrix of the climatic anomalies in Fig. S2. To exclude the spatial variation in *α* and to focus on the temporal variation in *α*, we deployed our aforementioned ridge regression model on annual mean climate per site to obtain an estimated mean *α* (α̂) per site and then calculated the difference between measured *α* and α̂ per site-year as *α*_rsd_. *α*_rsd_ indicates the variations that could not be explained by the spatial variability in climate variables but could be attributed to climate interannual variability.

We then explained the dependence of *α*_rsd_ on climatic anomalies and biome information using two machine learning approaches, random forest and XGBoost, a supervised machine learning algorithm with a gradient boosting framework (Chen & Guestrin, 2016), which both have been widely used in non-linear regression analyses (Janssen et al., 2023; Y. Yan et al., 2024). We determined the best hyperparameters in the random forest and XGBoost models with a grid search method and two-fold cross-validation using R ‘*caret*’ (Kuhn, 2008), ‘*randomForest*’ (Liaw & Wiener, 2002), and ‘*xgboost*’ (Chen & Guestrin, 2016) packages and extracted the relative importance of variables for the prediction. Our best model (random forest) explained 55.4% of the variation in *α*_rsd_. We ran the random forest model 50 times and interpreted the relative significance of each climatic anomalies using the percentage of explained mean squared error (%MSE, individual divided by the sum) and the partial effects of climatic anomalies on *α*_rsd_ with Shapley Additive exPlanations (SHAP) values which were extracted with the ‘*fastshap*’ package (Greenwell, 2024).

## Results

### Spatial variation in *α* and its environmental drivers

We noticed a strong spatial variation in *α* not only among biomes (> 10-fold) but also within each biome (Fig. 2a). Specifically, we found high average *α* values from forest (DBF: 0.043 mol CO_2_ mol photons^−1^, EBF: 0.037 mol CO_2_ mol photons^−1^, ENF: 0.034 mol CO_2_ mol photons^−1^, and MF: 0.034 mol CO_2_ mol photons^−1^), crop (CRO: 0.037 mol CO_2_ mol photons^−1^), and grassland (GRA: 0.035 mol CO_2_ mol photons^−1^) sites, and low values from savanna (SAV: 0.015 mol CO_2_ mol photons^−1^), shrubland (SH: 0.011 mol CO_2_ mol photons^−1^), and wetland (WET: 0.025 mol CO_2_ mol photons^−1^) sites (Fig. 2a). Using ridge regression and *lasso*, we examined the drivers of the spatial variations in *α* and found that water availability (SWC and VPD) was the largest driver (Fig. 2b, e, f). With a 100% increase in SWC and VPD, *α* increased by 29.8% and decreased by 28.0%, respectively (Fig. 2b). This was confirmed by the negative correlation between *α* and log_10_(AI) (*R* = −0.63, Fig. S3), which suggested a lower *α* in arid ecosystems. The LAI was found to be the second largest driver for the spatial variation in *α*. With a 100% increase in LAI, *α* increased 21.2% (Fig. 2b, h). In contrast, CO_2_ and DF played less significant roles with only 6% and 9.8% increases in *α* when CO_2_ and DF increased 100% respectively (Fig. 2b, d, g). Overall, TA positively impacts *α* (15% increase in α with a 100% increase in TA, Fig. 2b); however, we found that this impact turned negative at sites with high VPD (Fig. 2c).

**Fig. 2.**
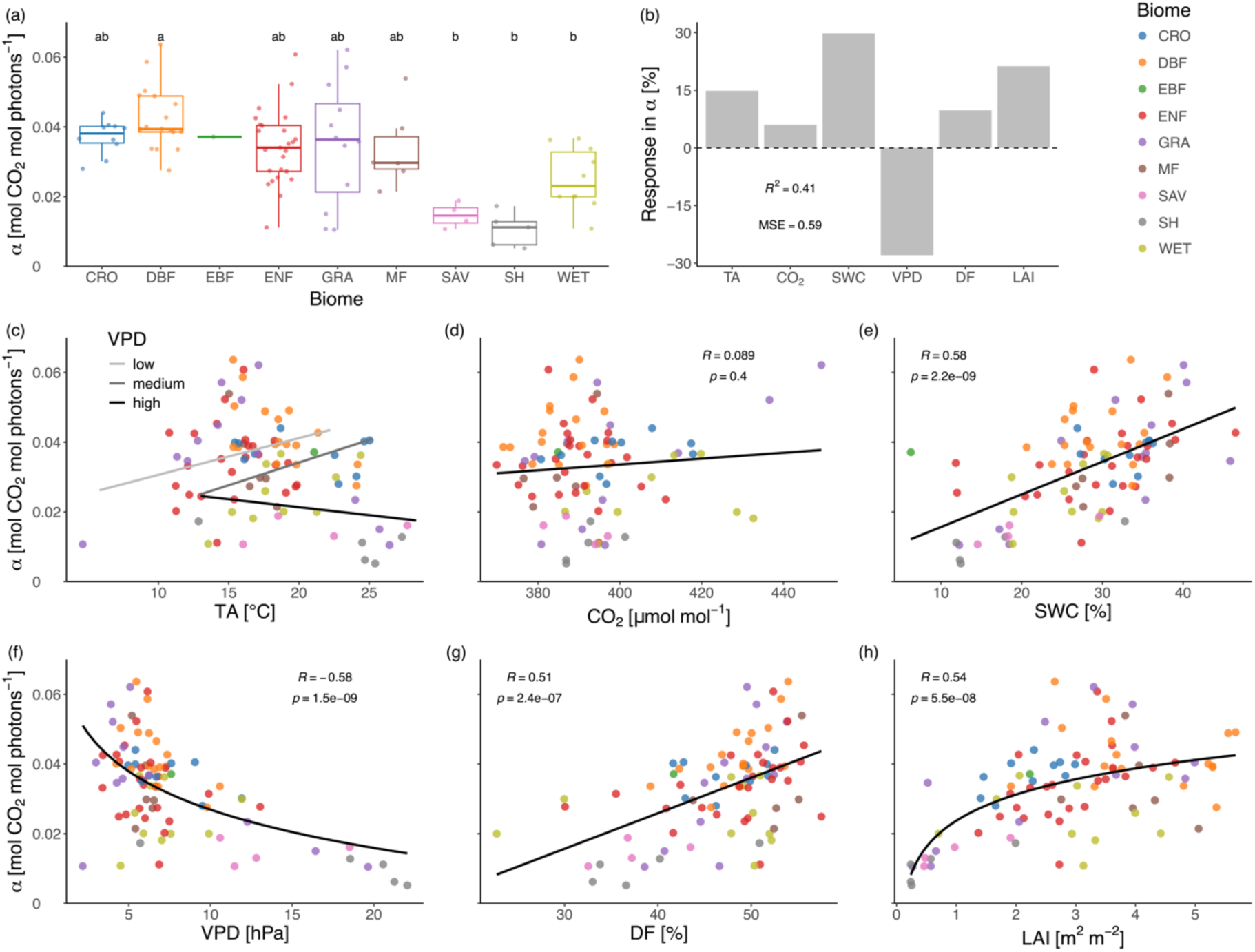
Spatial variations in photosynthetic quantum yield (*α*) and the dependence of the variations. (a) *α* was derived from eddy-covariance observations. The biomes we studied include CRO (croplands), DBF (deciduous broadleaf forests), EBF (evergreen broadleaf forests), ENF (evergreen needleleaf forests), GRA (grasslands), MF (mixed forests), SAV (savannas), SH (shrublands), and WET (wetlands). Each data point indicates a site-mean (growing season) *α* across all years. Different letters above bars in (a) indicate significantly different means of *α* (*p* < 0.05, analysis of variance and multi-comparison with Tukey’s HSD test), while different colors of the data points indicate biomes. We did not apply multi-comparison to EBF (*n* = 1). (b) Mean coefficients of variables (air temperature, TA; root zone soil water content, SWC; air vapor pressure deficit, VPD; diffuse light fraction, DF; and leaf area index, LAI), coefficient of determination (*R*^2^), and mean squared error (MSE) from the ridge regression and *lasso* models that estimate *α*. All variables including *α* were standardized prior to the analysis and a coefficient of 1 for a variable indicates that *α* increases 100% with a 100% change in the level of that variable. (c–h) Single-factor linear regression of *α* on independent variables. Different colors of the data points indicate biomes (see the legend in the top right). Regression lines (c–e, g: linear; f, h: logarithmic), *R*^2^, and *p* values are shown. In (c), data were grouped upon three VPD levels (low, medium, and high) before regression.

### Temporal trend in *α*

We examined the trend of *α* at each site and found only 13.3% (12 out of 90) of the sites showed a significantly positive trend (*p* < 0.05, Fig. 3). Most of sites with significantly positive trend in *α* were ENF and GRA: five out of 25 ENF and four out of 12 GRA sites showed a significantly increasing *α* (Fig. 3d, e), while only two ENF and one GRA sites had a decreasing *α*, respectively. Two out of six MF sites showed positive trend with one showing negative (Fig. 3f). Meanwhile, three out of 17 DBF sites showed a significant decrease in *α,* whereas only one site showed an increase (Fig. 3b). Other biomes (CRO, EBF, SAV, SH, and WET) did not show any significant trend in *α* over the years (Fig. 3a, c, g–i). Considering the length of the eddy covariance records, the trend detection is still subject to the strong interannual variations of variables.

**Fig. 3.**
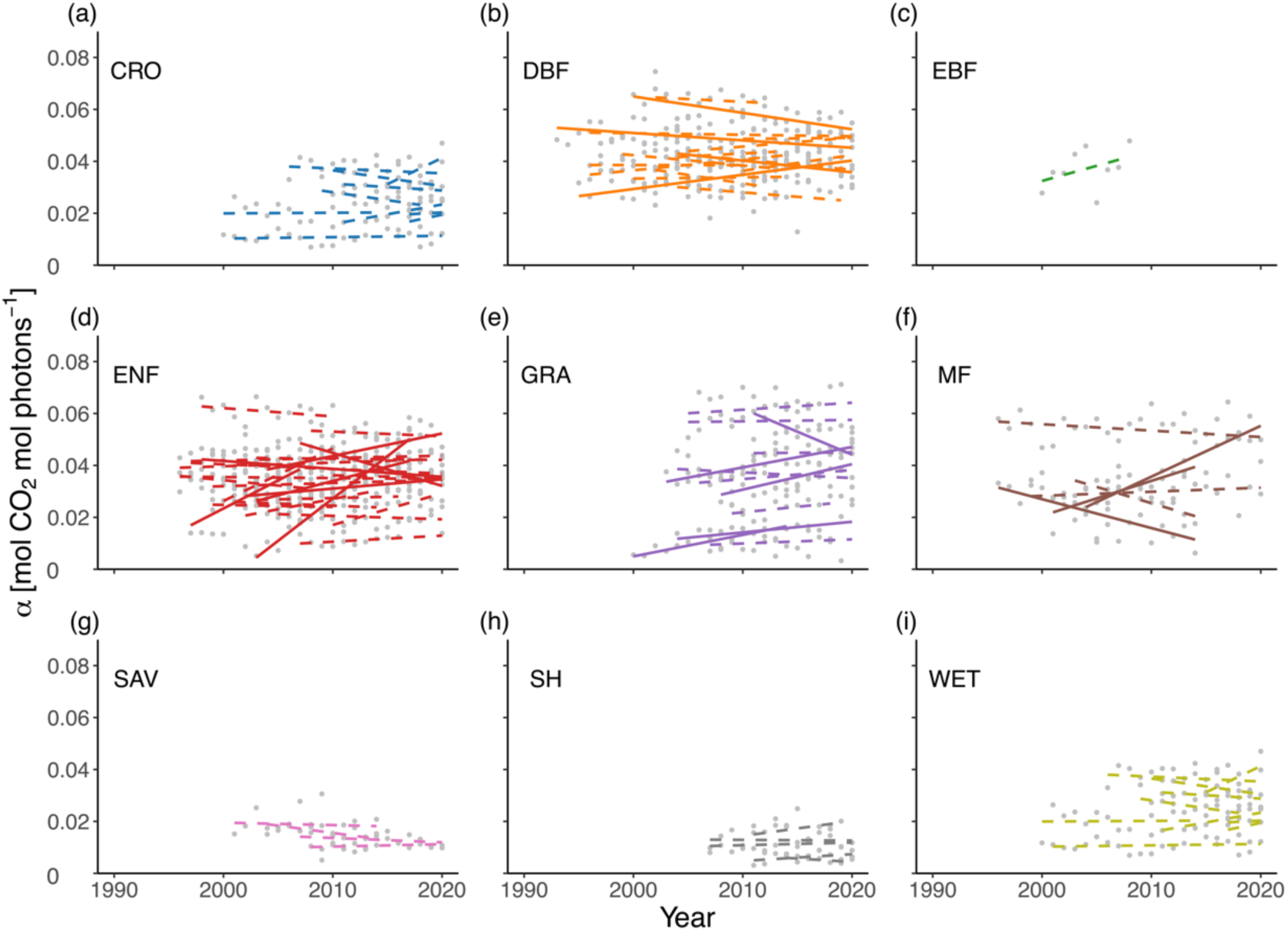
Trends of *α* over the years for each of the sites. Sites were grouped in biomes. Each grey dot indicates one site-year observation of growing season *α*. Solid lines indicate the site with a significant trend of *α* over the years (*p* < 0.05) while dashed lines indicate no significant trends.

### Temporal variation in *α* and its environmental drivers

We further examined the interannual variability of *α* over years for each site using coefficient of variation (CV). We found an average CV of 19.78% across all sites (Fig. 4a). While most biomes showed CV of *α* around 20%, we found the average CV for MF (27.03%) and SH (28.30%) were considerably greater than the average, and that of DBF (12.94%) were considerably smaller than average. Those indicate that MF and SH are likely to have larger interannual changes in *α*. We then quantified the contribution of climatic anomalies to the temporal variations in *α* (*α*_rsd_) using a random forest model (see Methods). We found that VPD_anom_ contributed the most (41.87%) to *α*_rsd_, followed by CO_2anom_ (13.16%), TA_anom_ (12.66%), and LAI_anom_ (12.19%), whereas SWC_anom_ (10.63%) and DF_anom_ (9.49%) played minor roles (Fig. 4b). We evaluated the partial effect of each climatic anomalies on *α*_rsd_ using SHAP values and found that VPD_anom_ negatively and linearly impacted *α*_rsd_ (Fig. 4f) while CO_2anom_ clearly had a positive impact on *α*_rsd_ (Fig. 4d) Meanwhile, we observed an increase in *α*_rsd_ with increasing LAI_anom_ but such effect saturated under positive LAI_anom_ (Fig. 4h). We did not find obvious response of *α*_rsd_ to TA_anom_, SWC_anom_, and DF_anom_.

**Fig. 4.**
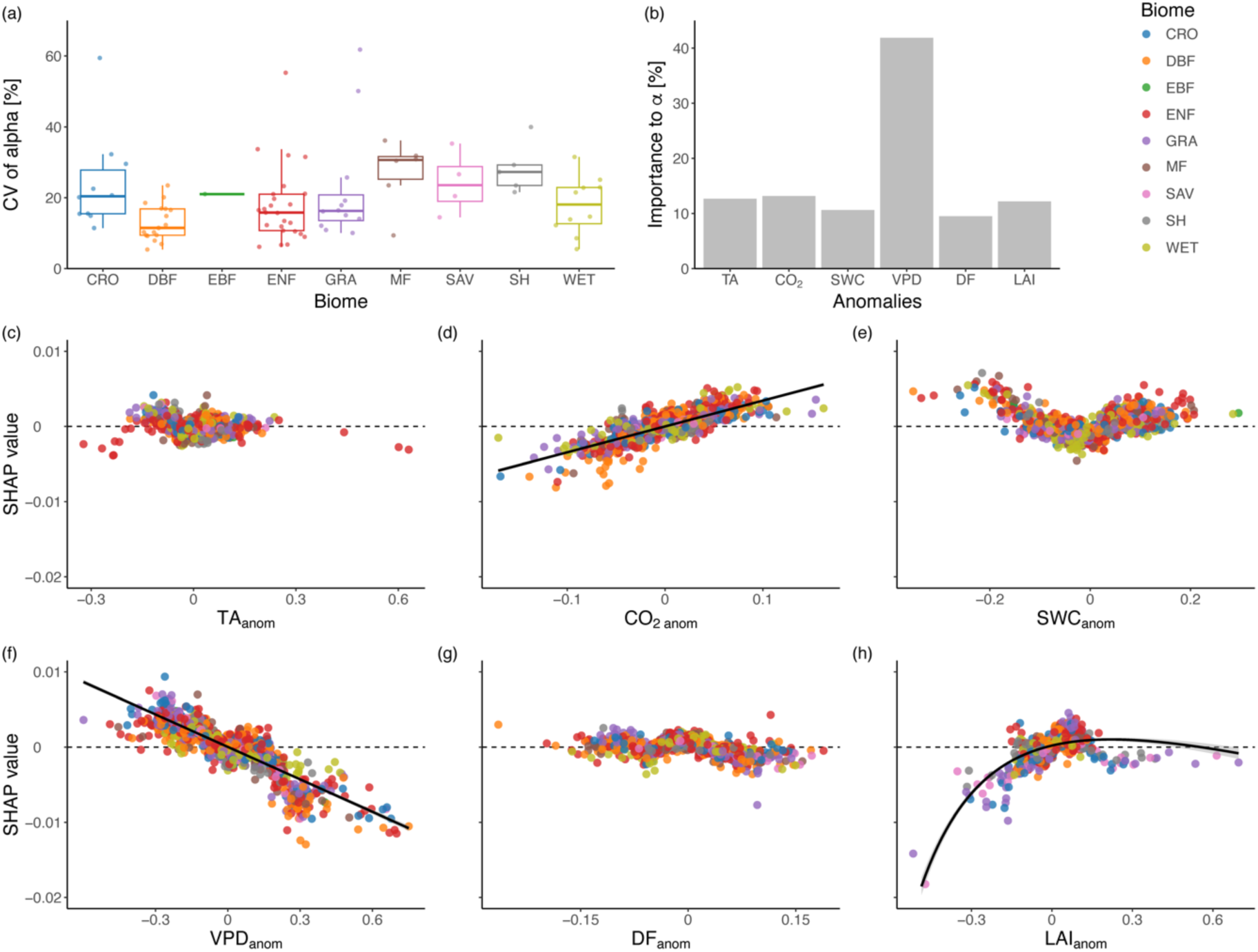
Interannual variability of *α* (a), relative importance of climatic anomalies to explaining the temporal variations in *α* (*α*_rsd_) using a random forest model (b), and Shapley Additive exPlanations (SHAP) values of each climatic anomalies to *α*_rsd_ from the model (c–g). In (a), we determined the coefficient of variation (CV) of the time series of annual-average *α* across sites. In (b), *α*_rsd_ denotes the difference between the observed annual *α* and the estimated *α* using the ridge regression model and observed annual climate (see Methods). Positive SHAP values indicate a positive effect on *α*_rsd_. The palette of data points in (b–g) indicates biomes (see the legend in the top right). Black solid lines in (b, e, and g) indicate the relationship between climatic anomalies and their SHAP values to *α*_rsd_ (b and e: linear; g: saturating).

### Comparison between LUE and *α*

The spatial variation in LUE estimated from the eddy covariance observations exhibited a pattern similar to that of *α*, though *α* is only one of the factors influence LUE. Specifically, CRO and forested sites had higher LUE values (0.0299 and 0.0256 mol CO_2_ mol photons^−1^ respectively) than SAV, SH, and WET (0.173, 0.131, and 0.197 mol CO_2_ mol photons^−1^ respectively, Fig. 5a). Within the biomes, we also found large variations with > 2-fold differences in LUE. LUE and *α* were significantly correlated (*R*^2^ = 0.89, *p* < 0.001, Fig. 5b), and *α* was 39% larger than the overall LUE, reflecting that *α* is the maximum LUE. *α* also dominated the temporal variation in LUE for all study sites, with a median *R*^2^s > 0.90 for all biomes (Fig. 5c), indicating that the climate impacts on LUE were largely propagated from their impacts on *α*. We further investigated the influence of climatic variables and LAI on the spatial variations in LUE (Fig. S4). We found that LUE negatively correlated to VPD (*R* = −0.33) and positively to SWC (*R* = 0.4), whereas none of the influences of TA, CO_2_, CI, and LAI were significant (*p* > 0.05).

**Fig. 5.**
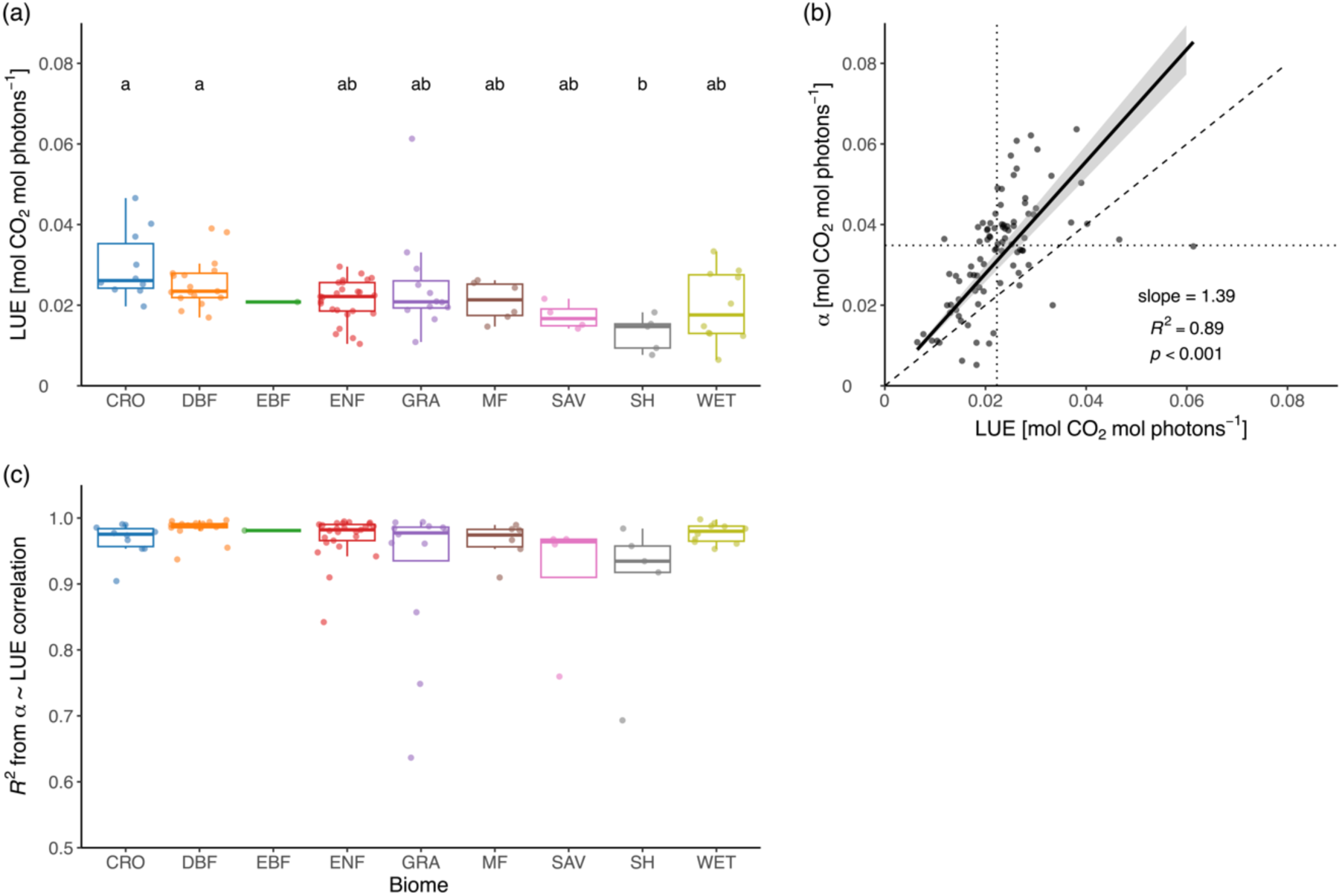
Variations in light use efficiency (LUE) and correlation between LUE and *α*. (a) LUE values of biomes. Different letters indicate significantly different levels of LUE (*p* < 0.05). We did not apply multi-comparison to EBF (*n*=1). (b) The linear relationship (*y* = *kx*) between site-mean *α* and LUE, with the regression line, confidence interval, *R*^2^, and *p-*value shown. The dotted lines indicate the medians of *α* and LUE. The dashed line indicates a 1:1 ratio of LUE to *α*. (c) The *R*^2^ from correlations (LUE *= kα*) between the time series of annual *α* and LUE at each site.

## Discussion

In this study, we investigated *α* from global long-term eddy covariance observations and examined the climatic drivers of spatiotemporal variations in *α*. We demonstrated that *α* varies largely among and within biomes, with higher values from CRO and forested sites than that from other biomes. We then identified water availability (VPD and SWC) as the dominant drivers of spatial variation in *α*, while VPD was the dominant constraining drivers of temporal variation in *α* which largely weakened the positive effect of CO_2_ and LAI. We also found that the variations in LUE were mainly explained by *α*, therefore the dependence of LUE was largely propagated from the climate dependence of *α*. Here, we discuss the mechanisms underlying the environmental regulation of *α* and provide insights into how to incorporate this information into ecosystem photosynthesis simulations.

### Biome dependence and spatial variation of *α* and climatic drivers

Our study showed a large variation in *α* between biomes (Fig. 2a). The higher *α* values at CRO and forested sites and lower levels at WET and SH sites are consistent with previous studies (Gower et al., 1999; He et al., 2022; Kergoat et al., 2008; Madani et al., 2014; Sandoval et al., 2023; Schwalm et al., 2006; Wei et al., 2017). This biome-specific variation could be attributed to the higher LAI and more layered structure of the canopy at forested sites (Sandoval et al., 2023), the higher leaf nitrogen content and light conversion efficiency at CRO sites due to management (fertilization and irrigation) and breeding (Gower et al., 1999; Turner et al., 2003; X.-G. Zhu et al., 2010), and the lower LAI and water availability at SAV, SH, and WET sites (Fig. 1e&h). We also found a large variation in *α* within each biome, suggesting that the biome alone cannot explain *α* spatially and that the prescribed *α* values embedded in most LUE models are highly likely to vary across sites (Running et al., 2004; Y. Zhang et al., 2017). We found that *α* spatially depends on SWC, VPD, CI, and LAI. These results are consistent with some previous studies (Bloomfield et al., 2023; Reich et al., 2018; Sandoval et al., 2023) and emphasize the advantage of prescribing *α* based on site climate and vegetation status, instead of assigning empirical values to biomes.

### VPD and soil moisture control spatial variations in *α*

We found that water availability is the main controlling factor of how *α* varies spatially (Chasmer et al., 2008; Garbulsky et al., 2010; He et al., 2022; Li et al., 2008; S. Wang et al., 2023). Low soil moisture (Barr et al., 2007; Jonard et al., 2022; Li et al., 2008; Q. Zhang et al., 2017) and high VPD (Bloomfield et al., 2023; Li et al., 2008; Reitz et al., 2023) intensify plant hydraulic stress and stomatal closure, shedding limitation on photosynthesis and *α*. Global warming has caused continuously increasing VPD (Novick et al., 2024) and more extremes in soil moisture on the land surface (Samaniego et al., 2018; Sherwood & Fu, 2014). Although debates remain regarding whether soil moisture or VPD is the most important driver of ecosystem production efficiency (Fu et al., 2022; Lu et al., 2022), our study indicates that their importance is equivalent in shaping spatial divergence of *α*. With more uneven distribution of SWC but the universally increased VPD, we might see that future *α* exhibits larger spatial variation.

Though TA enhances *α* over space under low and medium VPD levels, we found their correlation turned negative under high VPD, exhibiting an overall bell-shape response of *α* to TA (Fig. 2c). Both the bell-shaped response (i.e., often associated with the reporting of an optimal temperature (T_opt_) for highest *α* or LUE) and the linearly positive response have been reported by previous studies (Bloomfield et al., 2023; Li et al., 2008; Reitz et al., 2023; Sandoval et al., 2023). We suggest that VPD is a key factor modulating how TA impacts on *α*, especially at arid ecosystems (Zhong et al., 2023), and that T_opt_ is more likely to be detected for regions often under water stresses (Tan et al., 2017). Temperature exerts direct and indirect impacts on photosynthesis at the same time (Lloyd & Farquhar, 2008): directly through the changes in the activity of photosynthetic enzymes and the rate of respiration and photorespiration (the latter two affect the net flux of photosynthesis), and indirectly through the depression of photosynthesis due to associated higher VPD and lower soil moisture. Slot et al. (2024) demonstrated that neglecting the VPD effect misattributes the thermal sensitivity of photosynthesis in tropical species and leads to a significantly shifted T_opt_, while some species do not exhibit a T_opt_. Xu et al. (2022) found that low *α* values at TA above an apparent T_opt_ are due to the dramatically decreasing SWC above a threshold. Our findings suggest that the independent effect of TA on *α* is positive in most cases except for water-limited conditions, which is consistent with previous studies (Rogers et al., 2019; Schwalm et al., 2006). This could also explain why we observed more sites in ENF and GRA with positive α trends, as most of these investigated sites are located in temperate Northern Hemisphere with lower annual TA (Fig. 2c) and no severe water limitation (lower VPD, Fig. 1f). Therefore, we consider that the intrinsic T_opt_ of *α* at these ecosystems has not been reached under current warming.

### Effects of DF and LAI in variations of *α*

In our study, we found that DF and LAI also play a role in the spatial variations of *α*. DF influences *α* by altering the radiation profile within the canopy. High DF often occurs with a lower total incident PAR and a higher proportion of PAR penetrating deeper canopy (Chasmer et al., 2008; Y. Chen et al., 2021; Gu et al., 2002; Jenkins et al., 2007; Neimane-Šroma et al., 2024; Norman & Arkebauer, 1991; Reitz et al., 2023; Schwalm et al., 2006; Turner et al., 2003; Urban et al., 2007; Q. Zhang et al., 2017), leading to a higher ecosystem *α* as more light is utilized by the ecosystem. Our study confirms the role of DF in modulating *α* and then GPP across global ecosystems. Meanwhile, LAI determined the number or the area of leaves in a canopy. Studies have found a positive correlation between LAI and *α* (Bloomfield et al., 2023; Li et al., 2008; Medlyn, 1998). We suggest this relationship is not only due to the increase in the amount of leaves to conduct photosynthesis per area, but also due to the larger proportion of light being absorbed by shaded leaves – which tends to have higher light use efficiency (Osmond et al., 1988; Poorter et al., 2019). Our results emphasize the importance of considering different light use strategies at different layers of canopy when studying ecosystem photosynthesis, which has been discussed in a few recent modelling works (Bao et al., 2022; Guan et al., 2021; He et al., 2013; H. Yan et al., 2017). Furthermore, we found that positive DF_anom_ did not enhance *α*_rsd_ (Fig. 4g), meaning that *α* is likely to respond to interannual variations in DF weakly. We also found *α*_rsd_ increased with LAI_anom_ and then saturated, meaning more leaves enhance ecosystem *α* but with a limit (Li et al., 2008; Medlyn, 1998). It also indicates that in the traditional framework of LUE GPP models, LAI impacts not only APAR but also *α*.

### Dominant role of VPD in the temporal variations in *α* offsets the CO_2_ fertilization effect on *α*

CO_2_ has been thought to enhance LUE directly due to increased carboxylation and reduced photorespiration (Smith et al., 2020). To the best of our knowledge, our study is the first observation evidence that reports CO_2_ fertilization effect on *α* at ecosystem scales (Fig. 4d). However, this positive effect was remarkably smaller than the negative effect of VPD (Fig. 4f), which explains why we did not observe a widespread increase in α across most of the sites. Our observation of the global *α* trend is discrepant with the widely reported increases of *α* or LUE from leaf- or individual-level CO_2_ enrichment in labs (Long, 1991; Osborne et al., 1997), canopy-scale studies with FACE systems or chambers (Ainsworth & Long, 2005; DeLucia et al., 2002; Hui et al., 2001; Hymus et al., 2003; Manderscheid et al., 2003; Monje & Bugbee, 1998; Norby et al., 2003). This could be also due to that the magnitude of CO_2_ elevation in the atmosphere (increased by *c.a*. 22% from 1980 to 2020, Dunn et al. (2022)) is much smaller than those studies (e.g., 263% in Monje & Bugbee (1998) and 87% in Luo et al. (2000)) and thus cannot offset the impact of rising VPD. Our study reinforces the significance of considering the impact of rising CO_2_ in LUE-GPP models but also emphasizes the necessity in addressing the interaction with dynamics of water availability (especially VPD) and canopy structure (LAI).

### Relations between *α* and LUE and climatic dependence of LUE

We found that spatial variations in LUE shared a similar pattern with *α*, with higher values at the forested and cropland sites (Fig. 2a, Fig. 5a). Variations in LUE could be largely explained by α (95.3 ± 3.3%) across all biomes (Fig. 5c). Although the remaining unexplained variations are likely due the stomatal responses to climate variation, our findings suggest that the variation in *α* could indicate the variation in LUE with high accuracy. This strong dependence of LUE on *α* is consistent with leave level evidence (Y. Wang et al., 2022) which found that the quantum yield of Photosystem II (proportional to leaf *α*, Björkman & Demmig (1987)) does not respond to changes in stomatal conductance (*G*_s_, often proportional to photosynthetic rate) until *G*_s_ gets extremely low (<0.1 mol H_2_O m^−2^ s^−1^). This tight coupling between *α* and LUE provide an explanation for the physiological basis underlying the robustness of LUE-GPP models.

### Concluding remarks

Using global long-term eddy covariance observations, we investigated the spatiotemporal variations in ecosystem *α* and the drivers for the variations. We found large spatial variations in *α* across and within biomes, and that water availability (VPD and SWC) is the dominant driver of the spatial variations in *α*, with lower *α* for arid ecosystems. Our findings also suggested that the temporal change in *α* is mostly caused by variation in VPD, meanwhile elevated CO_2_ and LAI also impact the temporal change in *α*. Our study highlights the dominant role of VPD in regulating spatial and temporal changes in ecosystem light use and provide insights for improving its representation in ecosystem photosynthesis models.

## Acknowledgments

All authors acknowledge the support from the NUS Presidential Young Professorship awarded to X.L. (A-0003625-03-00) and a Tier 2 Academic Research Fund from Singapore Ministry of Education (MOE-T2EP50222-0006). All authors would like to thank Dr. Yang Li for her help in editing and proofreading this manuscript.

## Author contributions

LY and XL designed the study. LY collected, analyzed, and visualized the data. XL secured the funding and resources for this study. RZ and TWS contributed to data analysis. LY wrote the original manuscript with significant revisions from XL, RZ, TWS, and JT.

## Data availability

This study used openly available eddy covariance measurements provided by the FLUXNET2015 Tier 1 dataset (https://fluxnet.fluxdata.org/data/fluxnet2015-dataset/), AmeriFlux FLUXNET Dataset (https://ameriflux.lbl.gov/data/download-data/), and ICOS release 2023-1 of Level 2 Ecosystem data (https://www.icos-cp.eu/data-products/ecosystem-release). Soil moisture and potential evapotranspiration data are available at the GLEAM portal (https://www.gleam.eu/#datasets). LAI data are available at GIMMS LAI4g v1.2 (https://zenodo.org/records/8035760).

## Code availability

All analyses were performed using R (v.4.4.1, R Core Team). We extracted *α* from eddy covariance data using the ‘*Reddyproc*’ package (Wutzler et al., 2018), and the customized code to extract *α* is available at https://github.com/fueco/GcAmaxVcmax/blob/main/Calculation_Amax.R. We extracted growing seasons using the ‘*phenofit*’ package and performed ridge regression using the ‘*glmnet*’ package, XGBoost using the ‘*caret*’ and ‘*xgboost*’ packages, and random forest using the ‘*caret*’ and ‘*randomForest*’ packages. The associated code will be provided publicly upon the acceptance of this manuscript.

## Supporting information

Additional supporting information can be found in a separate Supporting Information document.

