## Supplemental table and figures for "Vapor pressure deficit dominates the spatiotemporal variations in ecosystem photosynthetic quantum yield"

### Supporting information

**Table S1** Information of the 90 sites used in this study.

| Site code[a] | Latitude (°N) | Longitude (°E) | Biome type[b] | Data start (year) | Data end (year) |
|---|---|---|---|---|---|
| BE-Bra | 51.31 | 4.52 | MF | 1996 | 2020 |
| BE-Dor | 50.31 | 4.97 | GRA | 2011 | 2020 |
| BE-Lon | 50.55 | 4.75 | CRO | 2004 | 2020 |
| BE-Vie | 50.30 | 6.00 | MF | 1996 | 2020 |
| CA-Ca1 | 49.87 | −125.33 | ENF | 1996 | 2010 |
| CA-Ca2 | 49.87 | −125.29 | ENF | 1999 | 2010 |
| CA-Cbo | 44.32 | −79.93 | DBF | 1994 | 2020 |
| CA-Gro | 48.22 | −82.16 | MF | 2003 | 2014 |
| CA-LP1 | 55.11 | −122.84 | ENF | 2007 | 2020 |
| CA-Oas | 53.63 | −106.20 | DBF | 1996 | 2010 |
| CA-Obs | 53.99 | −105.12 | ENF | 1997 | 2010 |
| CA-TP1 | 42.66 | −80.56 | ENF | 2002 | 2017 |
| CA-TP3 | 42.71 | −80.35 | ENF | 2002 | 2014 |

| | | | | | |
|---|---|---|---|---|---|
| CA-TP4 | 42.71 | −80.36 | ENF | 2002 | 2014 |
| CH-Aws | 46.58 | 5.79 | GRA | 2006 | 2020 |
| CH-Cha | 47.21 | 8.41 | GRA | 2005 | 2020 |
| CH-Dav | 46.82 | 9.86 | ENF | 1997 | 2020 |
| CH-Fru | 47.12 | 8.54 | GRA | 2005 | 2020 |
| CH-Lae | 47.48 | 8.36 | MF | 2004 | 2020 |
| CH-Oe2 | 47.29 | 7.73 | CRO | 2004 | 2020 |
| CZ-BK1 | 49.50 | 18.54 | ENF | 2004 | 2020 |
| CZ-Stn | 49.04 | 17.97 | DBF | 2010 | 2020 |
| CZ-wet | 49.02 | 14.77 | WET | 2006 | 2020 |
| DE-Akm | 53.87 | 13.68 | WET | 2009 | 2020 |
| DE-Geb | 51.10 | 10.91 | CRO | 2001 | 2020 |
| DE-Gri | 50.95 | 13.51 | GRA | 2004 | 2020 |
| DE-Hai | 51.08 | 10.45 | DBF | 2000 | 2020 |
| DE-Hte | 54.21 | 12.18 | WET | 2009 | 2018 |
| DE-Hzd | 50.96 | 13.49 | DBF | 2010 | 2020 |
| DE-Kli | 50.89 | 13.52 | CRO | 2004 | 2020 |
| DE-Lnf | 51.33 | 10.37 | DBF | 2002 | 2012 |
| DE-Obe | 50.79 | 13.72 | ENF | 2008 | 2020 |
| DE-RuR | 50.62 | 6.30 | GRA | 2011 | 2020 |
| DE-Tha | 50.96 | 13.57 | ENF | 1996 | 2020 |
| DK-Sor | 55.49 | 11.64 | DBF | 1996 | 2020 |
| FI-Hyy | 61.85 | 24.29 | ENF | 1996 | 2020 |
| FI-Let | 60.64 | 23.96 | ENF | 2009 | 2020 |
| FI-Sod | 67.36 | 26.64 | ENF | 2001 | 2014 |
| FR-Fon | 48.48 | 2.78 | DBF | 2005 | 2020 |
| FR-Gri | 48.84 | 1.95 | CRO | 2004 | 2020 |
| FR-Lam | 43.50 | 1.24 | CRO | 2005 | 2020 |
| FR-LBr | 44.72 | −0.77 | ENF | 1996 | 2008 |

| | | | | | |
|---|---|---|---|---|---|
| GL-ZaH | 74.47 | −20.55 | GRA | 2000 | 2014 |
| IT-Col | 41.85 | 13.59 | DBF | 1996 | 2014 |
| IT-Cpz | 41.71 | 12.38 | EBF | 1997 | 2009 |
| IT-Lav | 45.96 | 11.28 | ENF | 2003 | 2020 |
| IT-MBo | 46.01 | 11.05 | GRA | 2003 | 2020 |
| IT-Ren | 46.59 | 11.43 | ENF | 1999 | 2020 |
| IT-Ro2 | 42.39 | 11.92 | DBF | 2002 | 2012 |
| IT-Tor | 45.84 | 7.58 | GRA | 2008 | 2020 |
| NL-Loo | 52.17 | 5.74 | ENF | 2008 | 2020 |
| RU-Fyo | 56.46 | 32.92 | ENF | 1998 | 2020 |
| SE-Deg | 64.18 | 19.56 | WET | 2001 | 2020 |
| US-ARM | 36.61 | −97.49 | CRO | 2003 | 2020 |
| US-Bar | 44.06 | −71.29 | DBF | 2004 | 2021 |
| US-Blo | 38.90 | −120.63 | ENF | 1997 | 2007 |
| US-BZB | 64.70 | −148.32 | WET | 2011 | 2021 |
| US-BZF | 64.70 | −148.31 | WET | 2011 | 2021 |
| US-BZS | 64.70 | −148.32 | ENF | 2010 | 2021 |
| US-EML | 63.88 | −149.25 | SH | 2008 | 2020 |
| US-GLE | 41.37 | −106.24 | ENF | 2005 | 2020 |
| US-Ha1 | 42.54 | −72.17 | DBF | 1991 | 2020 |
| US-Ho2 | 45.21 | −68.75 | ENF | 1999 | 2020 |
| US-Jo1 | 32.58 | −106.64 | SH | 2010 | 2020 |
| US-Jo2 | 32.58 | −106.60 | SH | 2010 | 2020 |
| US-KFS | 39.06 | −95.19 | GRA | 2007 | 2019 |
| US-Los | 46.08 | −89.98 | WET | 2000 | 2014 |
| US-Me2 | 44.45 | −121.56 | ENF | 2002 | 2020 |
| US-MOz | 38.74 | −92.20 | DBF | 2004 | 2019 |
| US-Mpj | 34.44 | −106.24 | SAV | 2008 | 2020 |
| US-Myb | 38.05 | −121.76 | WET | 2010 | 2021 |

| | | | | | |
|---|---|---|---|---|---|
| US-NC4 | 35.79 | −75.90 | WET | 2009 | 2021 |
| US-Ne2 | 41.16 | −96.47 | CRO | 2001 | 2013 |
| US-Ne3 | 41.18 | −96.44 | CRO | 2001 | 2013 |
| US-NR1 | 40.03 | −105.55 | ENF | 1998 | 2016 |
| US-Oho | 41.55 | −83.84 | DBF | 2004 | 2013 |
| US-PFa | 45.95 | −90.27 | MF | 1995 | 2014 |
| US-Ro1 | 44.71 | −93.09 | CRO | 2004 | 2016 |
| US-Seg | 34.36 | −106.70 | GRA | 2004 | 2021 |
| US-Ses | 34.33 | −106.74 | SH | 2007 | 2021 |
| US-SRM | 31.82 | −110.87 | SAV | 2004 | 2014 |
| US-Syv | 46.24 | −89.35 | MF | 2001 | 2014 |
| US-Ton | 38.43 | −120.97 | SAV | 2001 | 2014 |
| US-Tw1 | 38.11 | −121.65 | WET | 2011 | 2020 |
| US-UMB | 45.56 | −84.71 | DBF | 2000 | 2014 |
| US-UMd | 45.56 | −84.70 | DBF | 2007 | 2021 |
| US-WCr | 45.81 | −90.08 | DBF | 1999 | 2014 |
| US-Whs | 31.74 | −110.05 | SH | 2007 | 2020 |
| US-Wjs | 34.43 | −105.86 | SAV | 2007 | 2021 |
| US-Wkg | 31.74 | −109.94 | GRA | 2004 | 2021 |

[a]: see full site names and detailed information at https://www.icos-cp.eu/observations/station-network and https://ameriflux.lbl.gov/sites/site-search/.

[b]: CRO: cropland; DBF: deciduous broadleaf forest; EBF: evergreen broadleaf forest; ENF: evergreen needleleaf forest; GRA: grassland; MF: mixed forest; SAV: savanna; SH: shrubland; WET: wetland.

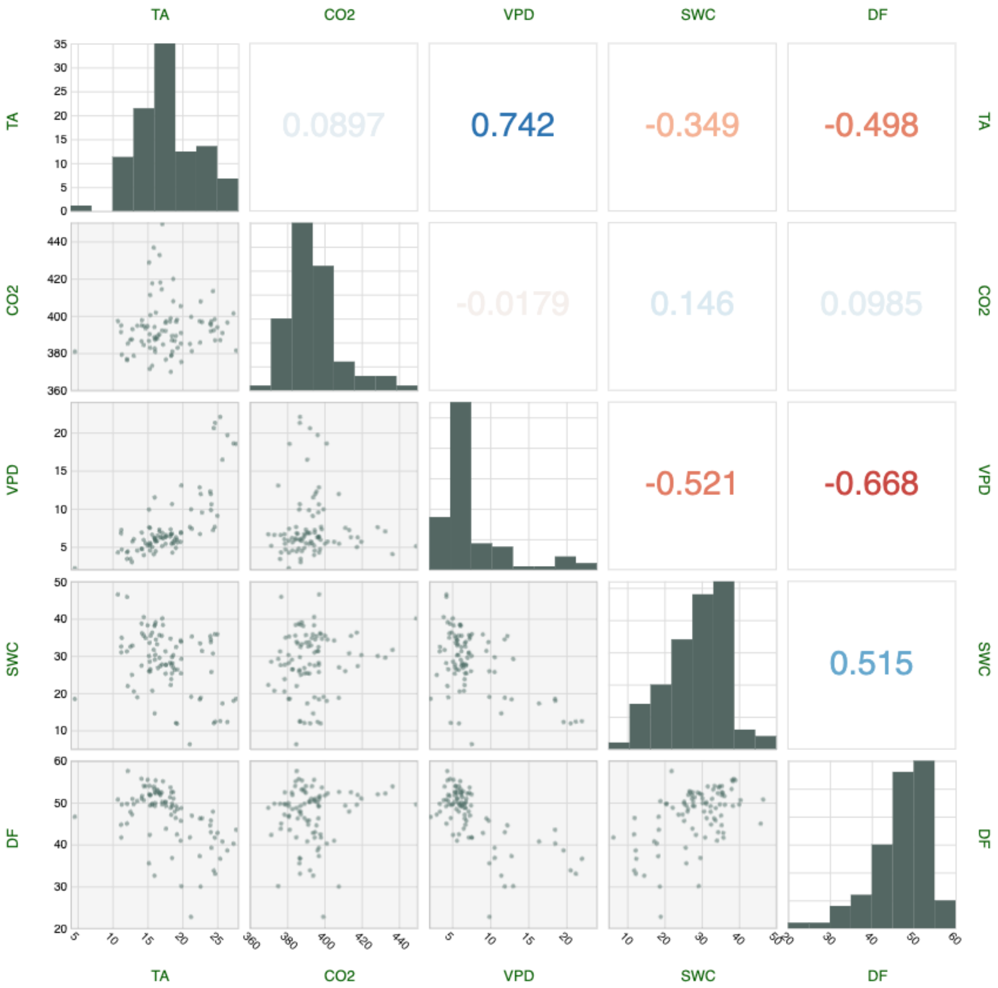


**Fig. S1** The correlation matrix of five climatic variables: air temperature (TA), ambient $CO_2$ concentration ($CO_2$), vapor pressure deficit (VPD), soil water content (SWC), and diffuse light fraction (DF). Values in the right-top panels indicate the coefficients of correlation.

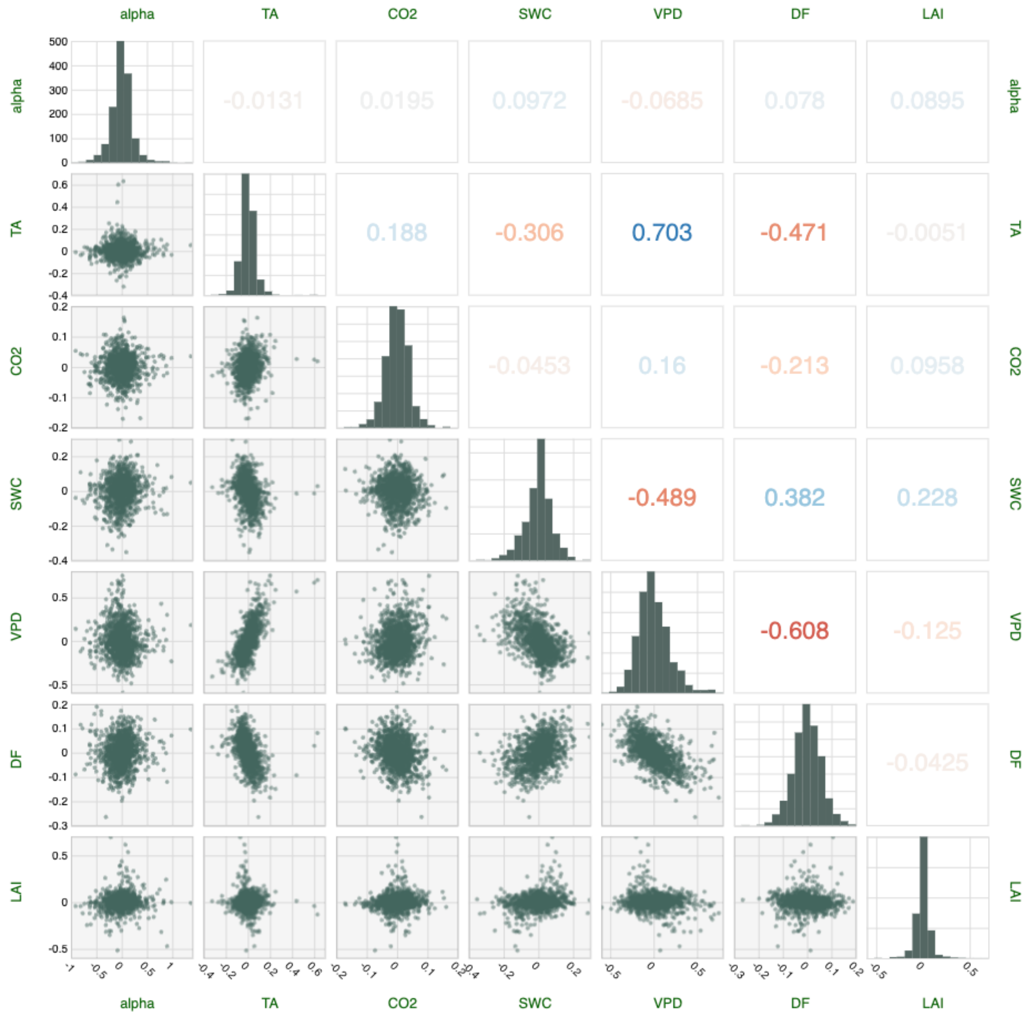


**Fig. S2** The correlation matrix of anomalies of quantum yield ($\alpha$), five climatic variables (TA, $CO_2$, VPD, SWC, and DF), and leaf area index (LAI). Values in the right-top panels indicate the coefficients of correlation.

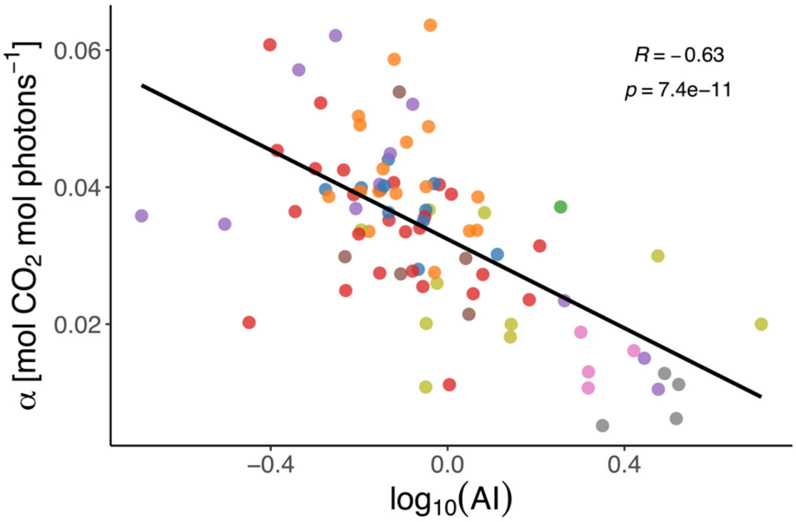


**Fig. S3** The correlation between aridity index (AI, logarithmic transformed) and $\alpha$. Higher AI indicates more arid conditions. Different colors of the data points indicate biomes the same as in Fig.1a. The linear regression line, $R$, and $p$ value are shown.

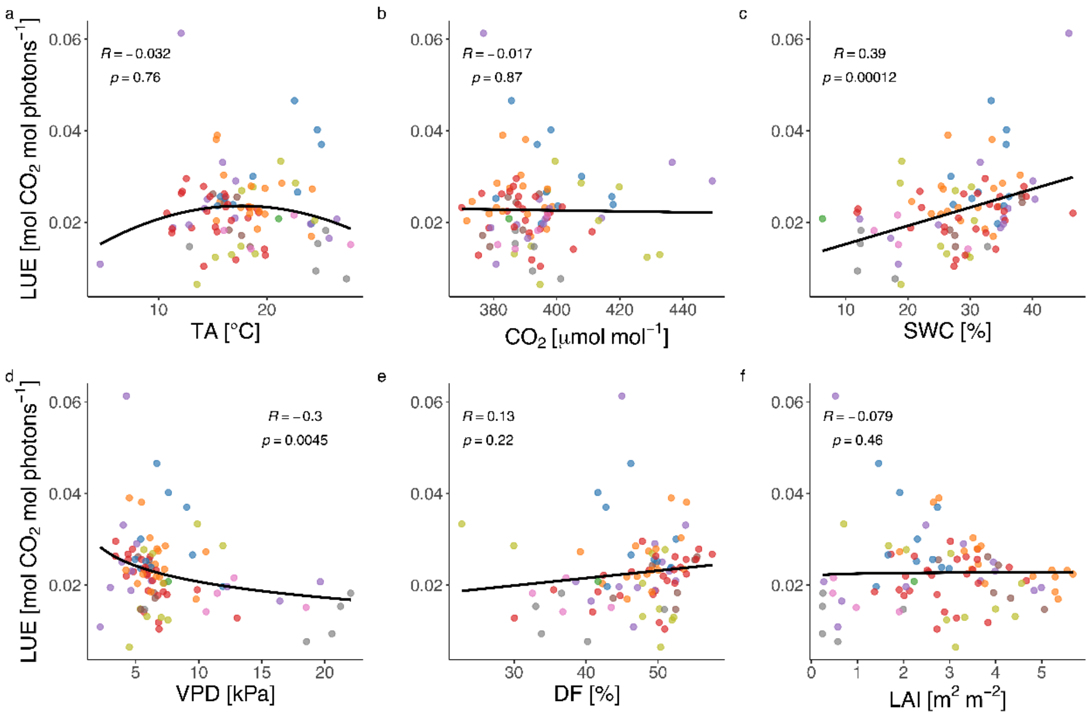


**Fig. S4** Spatial variations in light use efficiency (LUE) and the dependence of the variations on the variables: (a) TA, (b) $CO_2$, (c) SWC, (d) VPD, (e) DF, and (f) LAI. Different colors of the data points indicate biomes the same as in Fig.1a. Regression lines (b, c, e, f: linear; a, d: logarithmic), *R*, and *p* values are shown.
